# PERM1 Gene Delivery via AAV Prevents Heart Failure in a Mouse Model of Pressure Overload

**DOI:** 10.1101/2025.09.29.679055

**Authors:** Karthi Sreedevi, Ryan Montalvo, Abbigail Doku, Audrey Korte, Rebekah Thomas, Sarah Salama, Steven Burrows, Zhen Yan, Alexey V. Zaitsev, Junco S. Warren

## Abstract

Heart failure with reduced ejection fraction (HFrEF) remains a leading cause of mortality worldwide. A hallmark of HFrEF is impaired cardiomyocyte contractility accompanied by disrupted mitochondrial bioenergetics; however, no current therapy targets both pathologies simultaneously. PERM1, a striated muscle-specific regulator of mitochondrial bioenergetics, is downregulated in HFrEF patients. We recently demonstrated that overexpression of PERM1 via adeno-associated virus 9 (AAV9-PERM1) enhances both cardiac contractility and mitochondrial biogenesis in C57BL/6 mice. In this study, we evaluated the therapeutic potential of AAV9-PERM1 in a pressure overload-induced mouse model of HFrEF. C57BL/6 mice were treated with either AAV9-PERM1 or control AAV9-GFP (1×10^12^ GC/mouse), followed by transverse aortic constriction (TAC) surgery. At 4 weeks post-TAC, control mice receiving AAV-GFP exhibited reduced left ventricular ejection fraction (LVEF), whereas AAV-PERM1 preserved LVEF at baseline levels. This cardioprotective effect was sustained through 8 weeks. Notably, AAV9-PERM1 completely abrogated TAC-induced cardiac hypertrophy and fibrosis. Mitochondrial analysis revealed that AAV9-PERM1 preserved mitochondrial DNA copy number and TFAM protein levels, both of which were reduced by TAC in control hearts. AAV9-PERM1 also improved mitochondrial respiration using pyruvate and octanoylcarnitine as substrates and prevented TAC-induced impairments in oxidative capacity. While PGC-1α expression remained unchanged in control TAC hearts, it was modestly yet significantly upregulated by AAV9-PERM1 in both sham and TAC groups. In addition, AAV9-PERM1 suppressed TAC-induced increases in O-GlcNAcylation, a stress-related posttranslational modification of proteins. Co-immunoprecipitation further revealed interactions of PERM1 with creatine kinase and troponin C, key proteins in ATP transduction and contractility, suggesting a functional coupling between mitochondrial energetics and contractility. In conclusion, AAV-PERM1 gene therapy effectively preserves cardiac function under pressure overload by maintaining mitochondrial biogenesis, respiration capacity and contractility. This study further suggests AAV-PERM1 as a promising therapeutic strategy for HFrEF.

## 1. Introduction

Heart failure with reduced ejection fraction (HFrEF) is a life-threatening disease affecting about 27 million people worldwide and about 3 million people in the Unites States. The current treatment of HFrEF primarily focuses on neurohormonal antagonists, β-blockers, angiotensin-converting enzyme inhibitors (ACE), and angiotensin receptor blockers (ARBs)^1^. More recently, SGLT2 inhibitors, which was originally prescribed for type 2 diabetes, have been FDA-approved for both HFrEF and HFpEF patients. While these therapies have improved patient outcomes, high mortality and hospitalization rates persist, and current treatments do not target cardiomyocytes directly, limiting their ability to restore contractile function.

Emerging therapeutic strategy for HFrEF has been shifting toward approaches that directly improve the underlying defects in the heart rather than merely managing symptoms^2^. HFrEF hearts are known as an energy starved heart, reducing ATP contents by ~30% of healthy hearts^3^. Numerous studies demonstrated that mitochondrial impairment and dysfunction of energy metabolism make the heart vulnerable to pathological stress and contributes to the progression of HFrEF^4–6^. Over the past two decades, intensive studies have identified the transcription factors that orchestrate the expression of genes involved in mitochondrial bioenergetics in response to various physiological and pathophysiological stimuli^5–9^. Among these, PGC-1α is a key, nodal molecule in the maladaptive metabolic remodeling occurring in heart failure (HF), and its maintenance has been proposed as a potential therapeutic strategy to ameliorate mitochondrial impairment and contractile dysfunction^4^. However, studies have shown that transgenic (Tg) mice overexpressing PGC-1α developed cardiomyopathy due to uncontrolled mitochondrial biogenesis, which disrupts sarcomere integrity and contractile function^10^. In addition, moderate overexpression of PGC-1α (~3-fold) in mouse hearts accelerated the decline of systolic function and increased mortality during pressure overload despite the increase of mitochondrial gene expression and citrate synthase activity^11^. Similarly, overexpression and activation of key transcription factors regulating mitochondrial function and fatty acid metabolism, such as estrogen-related receptor (ERR) gamma and peroxisome proliferator-activated receptor (PPAR) in mice worsened cardiac dysfunction and reduced survival during pressure overload^12,13^. Thus, despite the recognized importance of maintaining cardiac energetics in pathological stress, there are currently no viable strategies targeting major metabolic regulators that can effectively preserve cardiac function. This may be due, at least in part, to a lack of coordination between the regulation of energetics and contractility under stress.

Efforts to enhance cardiac contractility for HFrEF therapy have long focused on the sarcomere, the fundamental contractile unit of muscle fibers, known as myofibrils, composed of thin actin and thick myosin filaments. Muscle contraction occurs when Ca^2+^ ions, released from the sarcoplasmic reticulum (SR), bind to troponin C (TnC) on the actin filaments, promoting steric movement of tropomyosin on actin that reveals binding sites for myosin heads, allowing the formation of cross-bridges. These cross-bridges then utilize ATP to cycle and generate force. Two classes of agents, inotropes and myotropes, aim to enhance cardiac contractility by either increasing the force of contraction or improving the efficiency of contractions, respectively. An issue with both inotropes and myotropes is that as they drive higher sarcomere contractility, it increases ATP utilization, which is often compromised in HFrEF^3^, thereby coming with an increased risk of worsening patient outcomes. Thus, despite on-going efforts to improve systolic function and energy metabolism, there are still no FDA-approved drugs that efficiently enhance cardiac contractility while preserving energy coupling in HFrEF.

PERM1, PPARGC1- and ESRR-induced regulator in muscle 1, is a striated muscle-specific regulator of mitochondrial bioenergetics^14^. We first identified PERM1 as a novel regulator of mitochondrial energetics as a downstream target of the striated muscle-specific epigenetic regulator SMYD1 in the heart, and showed that PERM1 is downregulated in HFrEF patients and in a mouse model of HFrEF^15^. In our follow-up study, we showed that *Perm1*-knockout (*Perm1*-KO) mice exhibit a moderate but statistically significant reduction in contractility and energy reserve in the heart, accompanied by global downregulation of proteins involved in oxidative phosphorylation (OXPHOS)^16^. Notably, unlike SMYD1, gene ablation of PERM1 was not embryonic lethal, and *Perm1*-KO mice exhibited a relatively mild phenotype. This distinguishes PERM1 from essential metabolic regulators such as PGC-1α and ERRα, suggesting its ability to fine-tune cardiac energy metabolism without disrupting homeostatic balance, and further supporting its therapeutic potential for HF. In this line of work, we generated adeno-associated virus (AAV) to perform gene delivery. Cardiac gene therapy via AAV has emerged as a promising option to treat advanced HF^17–21^. In this study, we demonstrate that gene delivery of PERM1 via AAV9-PERM1 to mouse hearts fully prevented the onset of HFrEF during pressure overload via transverse aortic constriction (TAC), a well-established mouse model of HFrEF. Our comprehensive mitochondrial bioenergetics assays further show that the cardioprotective effect of AAV9-PERM1 against pressure overload is correlated with fully preserved mitochondrial biogenesis and respiration capacity. These findings pave the way toward establishing a practical and translationally relevant approach for PERM1-based gene therapy in the treatment of cardiac disease.

## 2. Experimental procedures

### Animals and tissue harvest

All animal experiments were approved by the Institutional Animal Care and Use Committee (IACUC) at Fralin Biomedical Research Institute (FBRI) in Virginia Tech and were conducted according to the Guide for the Care and Use of Laboratory Animals. C57BL/6N female mice were purchased from Charles River. Animals were injected with AAV vectors carrying either gene of PERM1 or GFP (genetic control) at the age of 8-12 weeks. Four weeks after AAV vector injection, animals were subjected to TAC surgery. Hearts were harvested from these animals 8 weeks post-TAC surgery and were immediately frozen in liquid nitrogen and stored at −80 °C until they were used for analysis.

### AAV generation and administration in mice

The AAV9 vectors for PERM1 (AAV9.CMV.mPerm1-FLAG.hGH) and GFP (AAV9.CMV.PI.EGFP.WPRE.Bgh) were obtained from the Penn Vector Core (RRID: SCR_022432), University of Pennsylvania. AAV-PERM1 and AAV-GFP vectors were injected retro-orbitally into the venous sinus at a dose of 10^12^ genome copies (GC)/mouse, as previously performed ^22^.

### Echocardiography

Mice were subjected to echocardiography at four time points: before AAV injection, 4 weeks after AAV injection (immediately before TAC), and 4 and 8 weeks after TAC. All echocardiographic analysis were performed using Vevo 2100 (Visual Sonics) at FBRI. Mice were anesthetized in the induction chamber using 4% isoflurane in 100% O_2_ (2 L/min) and were maintained at 1.5 - 2.5% isoflurane and 0.8 L/min of O_2_ throughout the imaging studies. During the procedure, body temperature was maintained at 37.0 °C ± 1.0°C, and heart rate was maintained in the range of 500 ± 50 beats per minute. Initially, two-dimensional (2D) imaging (B-mode) was done to obtain a view along the parasternal short axis followed by M-mode imaging to obtain high-resolution measurements of left ventricular diameter and wall dimensions in systole and diastole. These measurements were used to estimate the left ventricular ejection fraction.

### Transverse Aortic Constriction Surgery

Transverse aortic constriction was performed on mice after 4 weeks of AAV-injection as previously described^23^. Briefly, mice were anesthetized with isoflurane (2% in O_2_) and surgery was performed. A single dose of Ethiqa (buprenorphine sustained release) (0.15 mg/kg) was administered subcutaneously before surgery. Mice were placed in a supine position and an incision of about 0.5cm length was made along the neck to expose aortic arch overlaying the trachea. The aortic arch was visualized under a low-powered microscope and a titanium micro clip (Horizon) was placed at the aortic arch between the innominate artery and the left common carotid artery using a ligation clip applicator that was calibrated to the diameter of 32-gauge size needle. Mice were sacrificed 8 weeks after TAC surgery to collect heart tissue. Control mice were subjected to sham procedure, in which all she steps of the standard surgical procedure were performed except application of the micro clip.

### Trichrome staining

The mouse hearts were embedded in the optimal cutting temperature (OCT) and cut into 10 µm sections. The sections were stained with Masson’s trichrome stain to evaluate cardiac fibrosis and imaged in the transmitted light at 10X magnification. The blue-green area occupied by collagen fibers in the interstitial space were measured using Image J software as described in detail in our previous publication [22] and presented as a percent of the total area.

### Western blotting analysis

Samples for Western blotting analysis was prepared and analyzed as we previously performed ^16^. Briefly, tissue samples were lysed in lysis buffer containing 50 mM Tris (pH 7.4), 10 mM EDTA, 1% sodium dodecyl sulphate (SDS), and protease inhibitors that include sodium butyrate (NaB), phenylmethyl sulphonyl fluoride (PMSF), sodium vanadate (Na3VO4), sodium fluoride (NaF) and EDTA-free protease inhibitor cocktail tablet. The lysate was centrifuged at 13,000 g for 10 min at 4°C and the supernatant was subjected to SDS electrophoresis followed by western blotting using 0.2 µm nitrocellulose membrane (Bio-Rad#1620112). The protein levels of PERM1, TFAM, PGC-1α, PGC-1β,ERRα,PPARα,NRF1,NRF2,Cytc, ATPsynthaseβ, CPT1b, CPT2, MCAD,GLUT4, OGT, OGA, OPA1,MFN1 were detected using rabbit antibodies (Sigma #HPA031711, Abcam #ab307302, Abcam #ab191838, Abcam #ab176328, Abcam #ab76228, Cayman product code #101710, Cell Signaling #D9K6P, Cell Signaling #D129C, Abcam #ab110325, Abcam #ab14730, Cell Signaling #E6M5M, Cell Signaling #E7D4W, Abcam #ab92461, Abcam #ab33780, Cell Signaling #24083, Abcam #ab124807, Abcam #ab56889-100 and Novus Biologicals #NBP2-34206) with 1:1000 dilution followed by goat anti-rabbit secondary antibody (Jackson ImmunoResearch Laboratories, Inc. #711-035-152) with 1:5000 dilution. The protein levels of total O-GlcNAcylated proteins and GLUT1 were detected using mouse antibodies (RL2, Abcam #ab2739 and Santa Cruz Biotechnology #sc377228) with 1:1000 dilution followed by anti-mouse secondary antibody from Abcam #ab6728 with 1:5000 dilution. The values of band intensities were normalized using β-tubulin (Abcam, #ab6046). The protein expression levels of β-tubulin were consistent in all groups that were used in this study. For the quantification of O-GlcNAc blots (Figure.5), the blots were divided into two portions: bands ≥75 kDa (High molecular weight referred as “HMW”) and bands <75 kDa (Low molecular weight referred to as “LMW”). In the HMW portion, non-specific bands around 75 kDa, which resulted from IgG band staining, were excluded by re-immunoblotting the same samples using anti-O-GlcNAc antibody pre-incubated overnight with 50 mM of an O-GlcNAc specific sugar, N-acetylglucosamine. Both HMW and LMW bands were quantified separately.

#### Mitochondrial Isolation

Cardiac muscle (~50-75mg) was separated for collection of a mitochondria-enriched fraction^24^ and placed immediately in 1mL of chilled mitochondria isolation buffer (MIB) [BSA (2mg/mL), Sucrose (70mM), Mannitol (210mM), HEPES (5mM), EGTA (1mM), pH 7.1]. At 4ºC, muscle was processed with a saw-tooth homogenizer for ~45 seconds. Homogenate underwent two centrifugations for 10 minutes at 4ºC at 800xg, then 9000xg. Following the second spin the cytosolic fraction was collected from the supernatant and immediately frozen at −80ºC. The final pellet was cleared of MIB by vacuum and resuspended in MIB free from BSA to be quantified by the Bradford method for normalization of protein loading for mitochondrial assays.

#### Mitochondrial Respiratory Flux (*J*O_2_)

##### Conceptual

Mitochondrial respiration was assessed using the creatine kinase clamp method ^25,26^ for two separate substrate conditions: (1) malate/pyruvate (M/P) and (2) malate/octanoylcarnitine (M/Oct). This method employs creatine kinase (CK) to maintain the stoichiometric ratio of ATP:ADP in a range that reflects *in vivo* conditions, increasing the physiological relevance of our findings above traditional experiments that utilize a supraphysiological bolus of ADP to assess maximal respiration. Further, this method allows for calculation of conductance by titration of phosphocreatine (PCr), which reflects a tractable bioenergetic stress test mimicking the transition from lower demand (ΔG_ATP_) to higher demand. Calculations of ΔG_ATP_ were performed using a freely available github calculator provided here (https://dmpio.github.io/bioenergetic-calculators/ck_clamp/) with conditions of 37ºC, 170mM ionic strength, 5mM creatine, 10mM phosphate, and pH 7.1 based on parameters referenced here (https://github.com/dmpio/bioenergetic-calculators/blob/master/jupyter_notebook/creatine-kinase-clamp.ipynb). Ranges of conductance included ΔG_ATP_ −12.94 kCal/mol for maximal respiration, −13.38, −13.71, −14.12, and −14.45 kCal/mol. Conductance of protons through the electron transport chain (reciprocal of resistance (1/R) is calculated as the slope over the linear range of *J*O_2_ vs. −13.38, −13.71, −14.12, and −14.45 based on Ohms law (I=V/R).

##### Procedural

Oroboros O2k (Innsbruck, Austria) was maintained at 37ºC and 500rpm in 0.5ml small volume chambers for each experiment. Chambers were calibrated with buffer D (BxD) [KMES (105mM), KCl (30mM), EGTA (1mM), KH_2_PO_4_ (10mM), MgCl_2_-6H2O (5mM), 0.05%BSA, pH7.1, solubility factor 0.966] to assess R1 (air saturation, ~200 µM O_2_) and R0 (zero oxygen) with sodium hydrosulfide (Sigma S1256). Respiratory assessments of oxygen flux (*J*O_2_ (pmol/sec/mg)) were made with BxD supplemented with 5mM creatine monohydrate and 30μg of mitochondrial protein. Following addition of mitochondria substrate-mediated respiration (non-phosphorylating) was performed for two separate substrate conditions: (1) malate (2.5mM) and pyruvate (5mM) and (2) malate (2.5mM) and octanoyl carnitine (0.2mM). Phosphorylating (maximal) respiration (ΔG_ATP_ −12.94) was assessed following the addition of ATP (5mM), creatine kinase (20mM), and PCr (1mM). Cytochrome c (0.005mM) was utilized to assess integrity with an *a priori* threshold of 15% increase in respiration, which did not require exclusion of any samples. Consecutive PCr titrations were then added at 1mM (ΔG_ATP_ −13.38 kCal/mol), 2mM (ΔG_ATP_ −13.71 kCal/mol), 4mM (ΔG_ATP_ −14.12 kCal/mol), and 6mM (ΔG_ATP_ −14.45 kCal/mol).

#### Mitochondrial *J*H_2_O_2_ Emission

##### Conceptual

Evaluation of oxidative stress and reactive oxygen species production was assessed via the Amplex Ultra Red (AUR)/ horseradish peroxidase (HRP) Assay^27^ under the identical creatine kinase conditions described above with some modifications. The AUR reaction is measures the production of H_2_O_2_ as it reacts with HRP to produce resorufin that is detected fluorometrically. Addition of superoxide dismutase (SOD) aids in the inner-mitochondrial conversion of superoxide radicals to H_2_O_2_, which is then emitted outside of the mitochondria to be detected by AUR. Further, a portion of superoxide could be buffered by endogenous antioxidants and limit clear interpretation of ROS production within muscle mitochondria. To this end, we included auranofin, a thioredoxin reductase inhibitor, as this is a primary buffering system within cardiac and skeletal muscle ^28^. Auranofin has been further demonstrated to not impact respiration or conductance ^25^. A standard curve was performed to convert resorufin fluorescence to H_2_O_2_ picomoles and *J*H_2_O_2_ (pmol/min/mg). Electron leak provides for an estimation of oxygen consumed by ROS that does not contribute to *J*O_2_ expressed as a percentage (*J*H_2_O_2_/ *J*O_2_ *100) after converting *J*O_2_ to pmol/min/mg at corresponding experimental conditions.

##### Procedural

Fluorescence was assessed via a fluorolog-QM (Horiba Scientific, Edison, NJ) outfitted with a 4-rotor turret system, temperature control, and stir bar capacity (Quantum Northwest, Liberty Lake, WA). Reactions occurred in a 600μl total volume with adapter cuvette at 37ºC, 500 rpm, 565:600 ex/em conditions over 4 minutes for each reaction to reach a steady state for each *J*H_2_O_2_ calculation. Buffer D described from *J*O_2_ experiments was supplemented further with SOD (20U/mL), HRP (1U/mL), auranofin (0.001mM), and AUR (0.02mM) and collected for a background rate. 600μg of mitochondria was added followed by substrates separately assessed under the M/P and M/Oct conditions for non-phosphorylating respiration (State 4 (S4). Phosphorylating respiration was assessed following addition of ATP (5mM), creatine kinase (20mM), and PCr (1mM). Conductance *J*H_2_O_2_ was assessed at ΔG_ATP_ −13.71 and −14.45 kCal/mol with 3mM and 10mM PCr additions, respectively.

**Mitochondria Isolation Buffer (pH 7.1)**
| **<u>Reagent</u>** | **<u>Source</u>** | **<u>Product #</u>** | **<u>CAS#</u>** |
| --- | --- | --- | --- |
| **Sucrose** | Fischer | BP220 | 57-50-1 |
| **Mannitol** | Sigma | M4125 | 69-65-8 |
| **EGTA** | Sigma | E4378 | 67-42-5 |
| **HEPES** | Gibco | 15630-080 | 7365-45-9 |
| **BSA** | Sigma | A6003 | 9048-46-8 |

**Buffer D (pH 7.1)**
| **<u>Reagent</u>** | **<u>Source</u>** | **<u>Product #</u>** | **<u>CAS#</u>** |
| --- | --- | --- | --- |
| **KMES** | Sigma | M0895 | 39946-25-3 |
| **KCl** | Sigma | P4504 | 7447-40-7 |
| **EGTA** | Sigma | E4378 | 67-42-5 |
| **KH<sub>2</sub>PO<sub>4</sub>** | Sigma | P0662 | 7778-77-0 |
| **MgCl<sub>2</sub>-6H<sub>2</sub>O** | Sigma | M2670 | 7791-18-6 |
| **BSA** | Sigma | A6003 | 9048-46-8 |

**JO<sub>2</sub> Mitochondrial Respiration Flux**
| **Reagent** | **Source** | **Product #** | **CAS#** |
| --- | --- | --- | --- |
| Sodium Hydrosulfide | Sigma | S1256 | 7775-14-6 |
| Potassium Pyruvate | Combi-Blocks | QA-1116 | 4151-33-1 |
| Malate | Sigma | M7397 | 97-67-6 |
| Octanoyl Carnitine | Sigma | 50892 | 25243-95-2 |
| Creatine Kinase from Rabbit muscle | Sigma | 10736988001 |  |
| Phosphocreatine | Sigma | P1937 | 108321-17-1 |
| ATP | Sigma | A2383 | 102047-34-7 |
| Cytochrome C | Sigma | C2506 | 9007-43-6 |
| Creatine Monohydrate | Sigma | C3630 | 6020-87-7 |

**JH<sub>2</sub>O<sub>2</sub> Amplex Ultra Red Assay**
| **Reagent** | **Source** | **Product #** | **CAS#** |
| --- | --- | --- | --- |
| Amplex Ultra Red | ThermoFischer | 36006 |  |
| Auranofin | Sigma | A6733 |  |
| Horseradish Peroxidase | Sigma | P8375 | 9003-99-0 |
| Superoxide Dismutase | Sigma | S9697 | 9054-89-1 |

### Mitochondrial DNA copy number measurements

DNA was extracted from heart tissues using the DNeasy Blood and Tissue kit (QIAGEN #69506) according to the manufacturer’s instructions. Mitochondrial copy number was measured as described in our previous publication ^22^. Briefly, 50 ng DNA was used to measure the relative copy number of mitochondrial DNA (mtDNA) and nuclear DNA (nDNA; for normalization) by qPCR using Taqman primers for mitochondrial gene ND1 and nuclear gene rpl32 respectively. Taqman primers used were as follows: for ND1: mt-ND1_CDU63R2, Rpl32: Mm07306626_gH.

### Gene expression analysis

Real-time PCR was performed using Taqman primers as previously described ^16^. Briefly, RNA was extracted using TRizol (Life Technologies), followed by ethanol extraction. Reverse transcription was performed from 1µg of RNA using QuantiTect Reverse Transcription Kit according to the manufacturer’s instruction (Qiagen). Taqman primers were purchased from ThermoFisher as follows: Nppb: Mm01255770_g1, Col3a2: Mm00802300_m1. The Cq values from the genes were normalized using Rpl32 (Mm07306626_gH).

#### Metabolomic Analysis

Untargeted metabolomic screening was performed using the Shimadzu GCMS-TQ8050 NX EI/CI/NCI mass spectrometry system, as described previously^16,29–31^. Briefly, freeze-clamped ventricle tissue samples (~12 mg) were homogenized in an extraction solution consisting of 90% methanol/water and containing d4-succinate and d4-myristic acid as internal standards. For plasma metabolomics, blood was collected using heparin at the time of terminal study and centrifuged at 1,500 x *g* for 10 minutes at 4°C. Ten microliters of plasma were mixed with 400 μL of the same extraction solution. All samples were dried and derivatized sequentially with O-methoxylamine hydrochloride in pyridine, followed by N-Trimethylsilyl-N-methyl trifluoroacetamide prior to GC-MS analysis. Bioinformatic analysis was conducted using MetaboAnalyst 6.0, which included the generation of heat maps and pathway impact analysis (**Figure 4**). Pathway analysis integrated p-values from pathway enrichment and visualized results where node color indicated statistical significance and node size reflected pathway impact values. Differences in metabolite abundance were evaluated using two-way ANOVA, with a p-value < 0.05 considered statistically significant. Data are presented as mean ± SEM.

### Statistical analysis

The data are presented as the mean ± standard error (SE), and all the data were analyzed using GraphPad Prism 10.0 software. Differences between the four groups were assessed using 2-way ANOVA. A value of p < 0.05 was considered statistically significant.

## 3. Results

### AAV-PERM1 prevents systolic dysfunction during pressure overload

Our previous study demonstrated that AAV9-PERM1 injection in healthy intact C57BL/6 mice achieved a three-fold increase in cardiac PERM1 expression which was associated with enhanced cardiac contractility measured 4 weeks post-injection^22^. To investigate whether gene delivery of PERM1 via AAV9-PERM1 can prevent the decline of systolic function caused by pressure overload, transverse aortic constriction (TAC) surgery was performed in C57BL/6 mice four weeks after retro-orbital administration of either AAV9-GFP (control) or AAV9-PERM1 (**Fig. 1A**). Hearts were harvested 8 weeks after TAC/Sham (12 weeks after AAV treatment) (**Fig. 1A**). Western blotting analysis confirmed our two previous findings (1) that PERM1 expression was significantly downregulated by TAC (61% of sham AAV-GFP, *p*<0.05) [15] and (2) that AAV9-PERM1 in healthy mice induces a strong increase in PERM1 expression [22] (**Fig. 1B-C**). To our great satisfaction, these two effects canceled each other, resulting in the maintenance of PERM1 at the normal level up to 8 weeks after TAC in mice treated with AAV-PERM1 prior to TAC. In other words, AAV-PERM1 *prevented* downregulation of PERM1 after TAC (**Fig. 1B-C**). Echocardiography was conducted at four time-points: (1) prior to AAV injection, (2) prior to TAC or sham surgery, (3) four weeks post-TAC, and (4) eight weeks post-TAC, just before terminal study (indicated by blue triangles in **Fig. 1A**). Consistent with our and other studies^16,32,33^, left ventricular ejection fraction (LVEF) was significantly reduced in AAV9-GFP mice after TAC (32% vs. 68% in GFP TAC vs. GFP Sham, p<0.001, **Fig. 1D-E**). Strikingly, AAV-PERM1 fully preserved LVEF at basal levels during pressure overload, with no significant difference observed between PERM1 Sham and PERM1 TAC groups up to 8 weeks post-TAC (64% vs. 66%, p>0.05, **Fig. 1D-E**).

**Figure 1.**
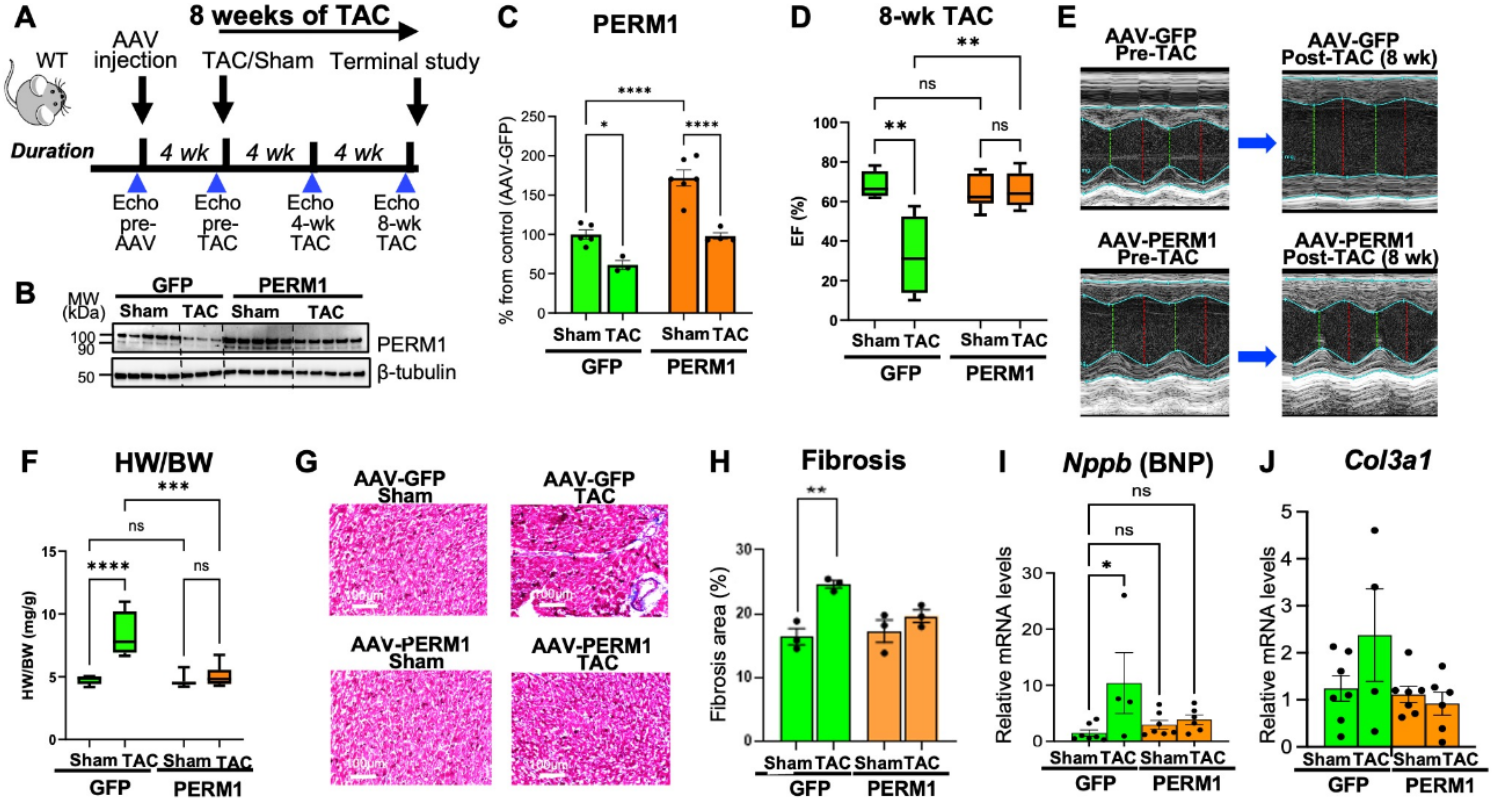
AAV-PERM1 prevents the major symptoms of heart failure during pressure overload in mice. **A**, Experimental design of the preventative study using AAV-PERM1 in mice. Blue triangles indicate time points at which echocardiography was performed. **B-C**, Western blot analysis of cardiac tissue showing that TAC-induced downregulation of PERM1 was normalized by AAV-PERM1. **D-E**, Left ventricular ejection fraction (LVEF) in Sham and TAC mice at 8 weeks post-TAC, along with representative M-mode echocardiographic images for each group. The decline in LVEF observed in TAC mice was fully prevented by AAV-PERM1. **F**, Heart weight-to-body weight (HW/BW) ratio was increased by TAC, while no significant difference was observed between AAV-GFP Sham and AAV-PERM1 TAC groups, indicating that AAV-PERM1 prevented pathological cardiac hypertrophy. **G-H**, Masson’s trichrome staining of cardiac tissue revealed increased fibrosis in TAC hearts, which was suppressed by AAV-PERM1. **I-J**, qPCR analysis of cardiac tissue showed that expression of *Nppb* (BNP), a marker of cardiac stress, was increased by TAC but was suppressed by AAV-PERM1 (**I**), whereas expression of *Col3a1*, a gene involved in collagen formation, was increased in a subset of AAV-GFP mice subjected to TAC, was not significantly affected overall (**J**). *: p<0.05, **:p<0.01,***:p<0.001, ****p<0.0001, ns: not significant, by two-way ANOVA.

TAC for 8 weeks significantly increased the heart weight to body weight (HW/BW) ratio in mice treated with AAV9-GFP (8.3 vs. 4.7 in GFP TAC vs. GFP Sham, p<0.05, **Fig. 1F**), indicating the development of cardiac hypertrophy. AAV-PERM1 administration abrogated TAC-induced pathological hypertrophy (5.1 vs. 4.6 in PERM1 TAC vs. PERM1 Sham, p>0.05, **Fig. 1F**). Masson’s trichrome staining of cardiac tissue revealed a significant increase in fibrosis following TAC in mice treated with AAV9-GFP (25% vs. 16.5% in GFP TAC vs. GFP Sham, p<0.05, **Fig. 1G-H**), whereas AAV-PERM1 treatment prior to TAC prevented this increase (**Fig. 1G-H**). Furthermore, qPCR analysis of cardiac tissue showed a significant increase in the mRNA levels of BNP (*Nppb*), a marker of cardiac stress, in GFP TAC hearts (10-fold increase in GFP TAC compared with GFP Sham, p<0.05), whereas BNP expression remained at basal levels in both PERM1 Sham and PERM1 TAC hearts (p>0.05 vs. GFP Sham, **Fig. 1I**). On average, the expression of *Col3a1*, a gene involved in collagen synthesis and fibrosis, was increased in GFP-TAC hearts (2.3-fold change vs. GFP Sham), but due to large dispersion of outcomes the difference did not reach statistically significant. Nevertheless, the excessively high levels of *Col3a1* observed in a subgroup of GFP-TAC hearts was not present in the PERM1 TAC group (**Fig. 1J**)

In summary, these results suggest that gene delivery of PERM1 completely prevents the three major pathological symptoms of heart failure: decline in the systolic contractile function, pathological hypertrophy, and fibrosis.

### PERM1 overexpression protects mitochondrial respiration and oxidative stress against TAC in a substrate-dependent manner

We previously demonstrated that adenovirus-mediated PERM1 overexpression enhances both glucose and fatty acid oxidative capacity in cardiomyocytes^15,36^. Mitochondrial respiratory capacity is known to be reduced in the failing heart, partly due to impaired fatty acid oxidation^37^. To investigate whether AAV-mediated gene delivery of PERM1 can preserve mitochondrial energetics under pressure overload, we performed a comprehensive analysis of mitochondrial respiration in isolated heart mitochondria from mice treated with either GFP or AAV-PERM1 and subjected either to Sham or TAC surgery (8 weeks). Mitochondrial respiration was assessed using the Oroboros O2k system. We evaluated Complex I-mediated respiration through the TCA cycle using pyruvate and malate (P/M), and fatty acid-dependent respiration using medium-chain fatty acid octanoyl-carnitine and malate (Oct/M). The creatine kinase clamp was employed to assess mitochondrial respiratory flux (*J*O_2_) under a range of physiological energetic demands by titrating phosphocreatine.

Under P/M-supported conditions, non-phosphorylating respiration was significantly reduced in TAC hearts compared to Sham in the GFP group (1066 vs. 353 pmol/s/mg, p<0.05). AAV-PERM1 administration significantly increased basal non-phosphorylating respiration compared to GFP Sham controls (1685 vs. 1066 pmol/s/mg, p<0.05; **Fig. 2A**). Remarkably, AAV-PERM1 preserved mitochondrial respiration under pressure overload, with PERM1-TAC values remaining comparable to PERM1-Sham levels (1440 vs. 1685 pmol/s/mg, **Fig. 2A**). A similar protective effect was observed in maximally stimulated respiration at ΔG_ATP_ −12.94 (**Fig.2B**). The protective role of PERM1 was further confirmed by an increased respiratory conductance (slope of *J*O_2_ vs. ΔG_ATP_) and improved respiration across all levels of energetic demand (**Fig. 2C-D**).

**Figure 2.**
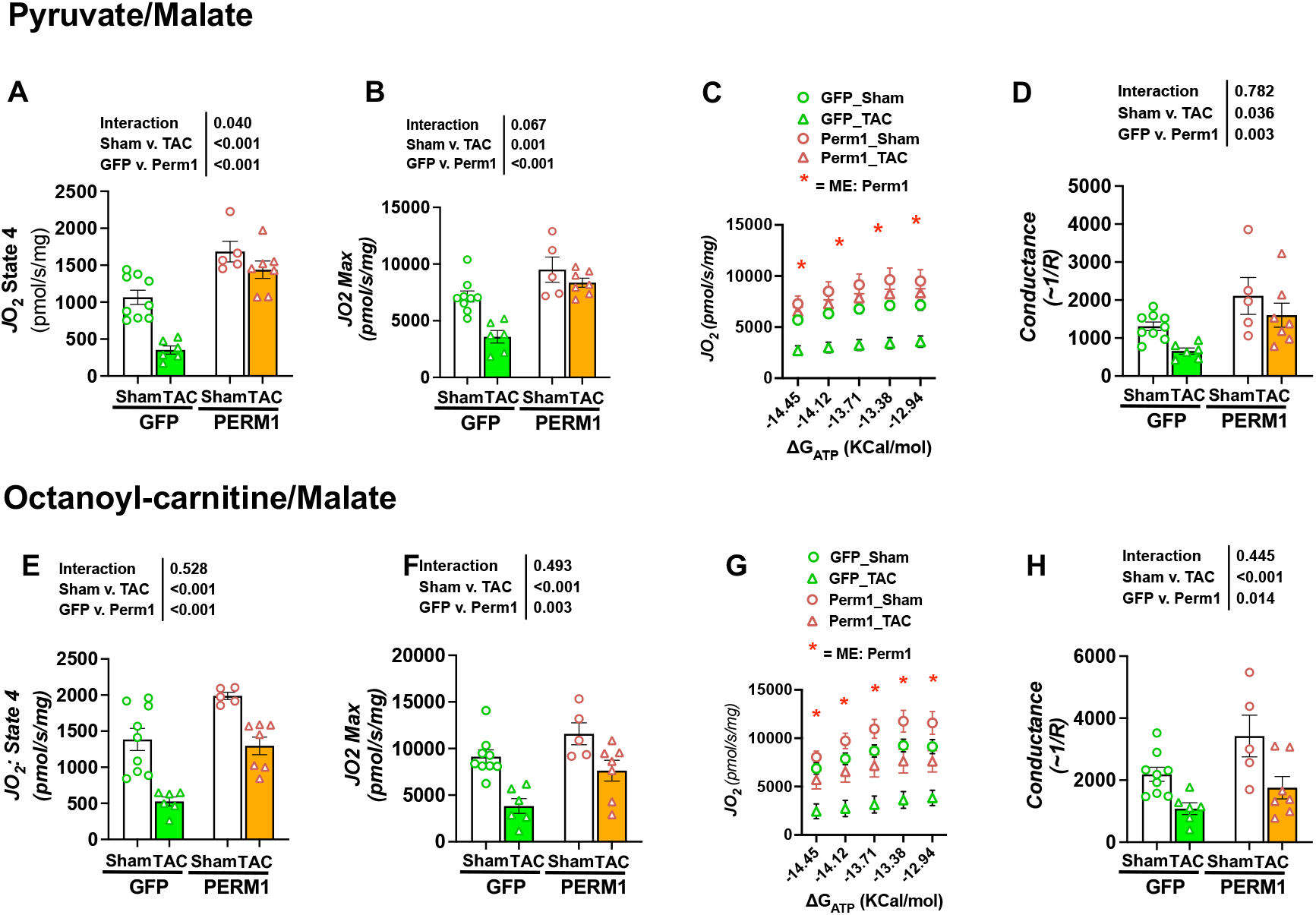
AAV-PERM1 counteracts TAC-induced depression of mitochondrial respiration in a substrate-dependent manner. Mitochondria were isolated from heart tissue of AAV-GFP and AAV-PERM1 treated mice 8 weeks after TAC or Sham surgery. Pyruvate/malate (**A-D**) or octanoyl-carnitine/malate (**E-H**) were used as mitochondrial substrates. For each substrate regimen we present, from left to right, JO2 in State 4 (**A** and **E**), maximal JO2 (at ΔG_ATP_ = −12.94 Kcal/mol) (**B** and **F**), JO2 as a function of ΔG_ATP_ (**C** and **G**), and conductance (the slope of JO2 - G_ATP_ relationship in the linear range of ΔG_ATP_) (**D** and **H**). TAC strongly reduced JO2 under all tested conditions. Under pyruvate/malate conditions, AAV-PERM1 fully prevented JO2 depression caused by TAC at all levels of energetic demand. Under octanoyl-carnitine/malate conditions, the ability of AAV-PERM1 to prevent JO2 depression caused by TAC was less than under pyruvate/malate conditions, yet AAV-PERM1 significantly improved respiration at all levels of energetic demand. Data presented as mean ± SEM and analyzed by 2-way ANOVA. Asterisks in Panels **C** and **G** indicates statistically significant main effect of transgene (PERM1 vs. GFP).

Consistent with these findings, the PERM1-Sham group exhibited significantly elevated non-phosphorylating respiration in the Oct/M condition compared to GFP-Sham (p<0.05), indicating enhanced fatty acid oxidation (FAO) capacity. Importantly, AAV-PERM1 treatment preserved basal *J*O_2_ under pressure overload, maintaining values comparable to GFP-Sham (**Fig. 2E**). However, unlike the M/P condition, the protective effect of PERM1 on non-phosphorylating respiration under Oct/M was diminished from the elevated baseline observed in the Sham group (1989 vs. 1296 pmol/s/mg, PERM1-Sham vs. PERM1-TAC, p<0.05, **Fig. 2E**). Maximal respiration at ΔG_ATP_ −12.94 was enhanced by AAV-PERM1 and remained preserved under TAC stress (**Fig. 2F**). Interestingly, baseline conductance in the Oct/M condition was more strongly enhanced by PERM1 overexpression than in Pyr/M, yet TAC still imposed a significant negative effect (**Fig. 2G–H**).

Pathological stress in heart failure is accompanied by elevated oxidative stress and excessive mitochondrial reactive oxygen species (ROS) production^38^. To determine whether preserved mitochondrial respiration by AAV-PERM1 also suppresses ROS, we measured H_2_O_2_ flux (*J*H_2_O_2_) in the same heart mitochondria used for *J*O_2_ analysis. Two key parameters were assessed: (1) the absolute H_2_O_2_ flux, and (2) the % electron leak—defined as (*J*H_2_O_2_ / *J*O_2_) x 100—representing the fraction of electron flow diverted to ROS production rather than ATP generation. The creatine kinase clamp allowed parallel assessment under identical respiratory states.

In the M/P condition, the absolute *J*H_2_O_2_ was unchanged under State 4 (**Fig. S1A)** but was increased by TAC under phosphorylating conditions, and this increase was not prevented by AAV-PERM1 (**Fig. S1B-D**). A very similar pattern was observed in the Oct/M condition (**Fig. S1E-H**). Electron leak, considered a more physiologically relevant indicator of mitochondrial redox imbalance, was significantly increased in AAV-GFP TAC group under all tested conditions, and this increase was partially or fully abolished by AAV-PERM1 in substrate- and ΔG_ATP_-dependent manner (**Fig. S2**). Specifically, the protective effect of AAV-PERM1 was stronger under P/M than Oct/M and tended to decrease with increasing energy demand.

In summary, these results show that AAV-PERM1 enhances both pyruvate- and fatty acid-mediated mitochondrial respiration in the heart, and that it prevents TAC-induced respiratory impairment. The protection appears substrate-dependent: PERM1 fully preserved respiration in the pyruvate condition, whereas protection in the fatty acid condition was partial. While AAV-PERM1 did not significantly suppress H_2_O_2_ production, it effectively reduced electron leak, highlighting its protective role in maintaining mitochondrial fitness during pressure overload.

### AAV-PERM1 preserves mitochondrial biogenesis and the expression of nuclear receptors during pressure overload

Genes and proteins involved in mitochondrial oxidative phosphorylation (OXPHOS) are primarily regulated by nuclear receptors and transcriptional coactivators such as estrogen-related receptors (ERRs), PGC-1 coactivators, peroxisome proliferator-activated receptors (PPARs), and nuclear respiratory factors (NRFs), which orchestrate the expression of OXPHOS genes in response to increased energy demands^37^. In contrast, mitochondrial dysfunction in the failing heart is often associated with downregulation of these transcription factors and cofactors ^38^. Our group and others have previously shown that PERM1 regulates the expression of ERRs and PPARα, along with their downstream targets involved in the mitochondrial electron transport chain (ETC) and fatty acid oxidation (FAO) ^16^. To determine whether the preserved mitochondrial energetics observed in AAV-PERM1-treated hearts (**Fig. 2**) is associated with the expression of these transcriptional regulators, we performed Western blot analyses using cardiac tissue from Sham and TAC mice (8 weeks) treated with AAV-GFP or AAV-PERM1. We found that TAC led to significant downregulation of ERRα, PPARα, NRF1, and TFAM in AAV-GFP hearts, all of which were maintained at a normal (or above normal) levels in AAV-PERM1 mice (p<0.05, GFP-TAC vs. PERM1-TAC, **Fig. 3A-H**). Interestingly, TAC did not affect PGC-1α or NRF2 protein expression in GFP-treated hearts (p>0.05), whereas AAV-PERM1 significantly increased PGC-1α levels in both Sham and TAC conditions (133% and 132% of GFP Sham, respectively, p<0.05 for both).

**Figure 3.**
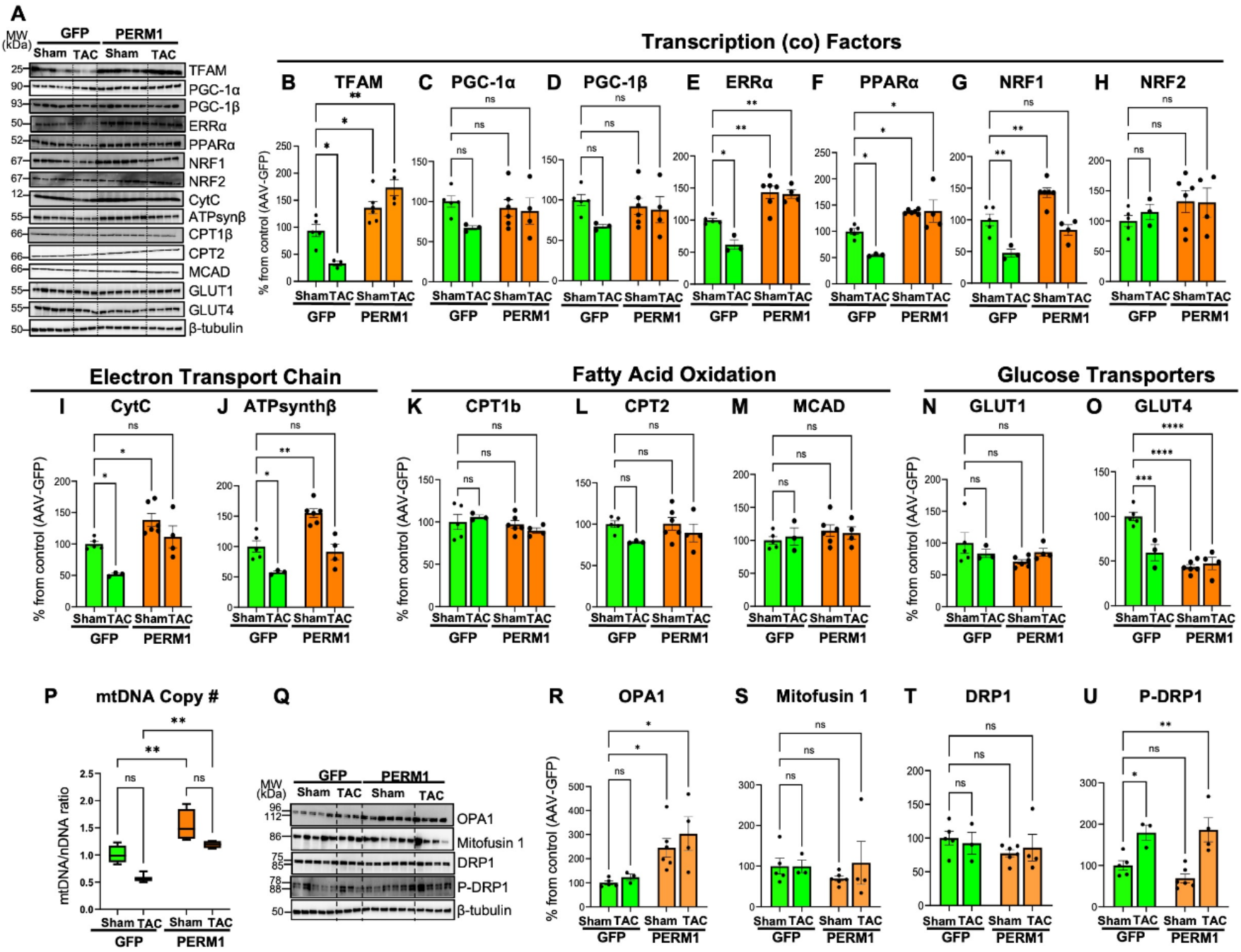
AAV-PERM1 preserves mitochondrial biogenesis and key regulators of mitochondrial bioenergetics during pressure overload. **A–O**, Western blot analysis of transcription factors, cofactors involved in mitochondrial bioenergetics, selected components of electron transport chain, selected key enzymes of fatty acid oxidation (FAO), and glucose transporters. (n=5 AAV-GFP-Sham, n=3 AAV-GFP-TAC, n=4 AAV-PERM1-Sham, n=6 AAV-PERM1-TAC). **P**, Quantification of mitochondrial DNA copy number showing a reduction in GFP TAC hearts, but full preservation by AAV-PERM1 treatment. (n=5 AAV-GFP-Sham, n=3 AAV-GFP-TAC, n=7 AAV-PERM1-Sham, n=4 AAV-PERM1-TAC). **Q–U**, Western blot analysis of proteins involved in mitophagy. Statistical comparisons presented are results of post-hoc pairwise Holm-Sidak test applied after two-way ANOVA. ****: p<0.0001, ***: p<0.001, *: p<0.05, mean ± SEM, ns: not significant

Consistent with the downregulation of ERRα and NRF1, their mitochondrial targets—cytochrome c (CytC) and ATP synthase β (Complex V)—were also reduced in GFP-TAC hearts (p<0.05 vs. GFP Sham), whereas in PERM1-TAC hearts the expression of these genes was maintained at least at the normal levels (p>0.05 vs. GFP Sham, **Fig. 3I-J**).

In contrast, neither TAC nor AAV-PERM1 affected the expression of CPT1b and CPT2 (the enzymes which mediate long-chain fatty acid uptake) and MCAD (enzyme responsible for metabolizing medium-chain fatty acids) (**Fig. 3K-M**). Lastly, myocardial protein expression of glucose transporter GLUT1 was not affected with either TAC or overexpression of PERM1, whereas the protein expression of GLUT4 was significantly decreased with AAV-PERM1 in both Sham and TAC hearts (43% and 47% of GFP Sham, respectively, p<0.05, **Fig. 3N-O**).

Mitochondrial biogenesis is impaired in the failing human heart due to reduced mitochondrial DNA (mtDNA) replication and mtDNA deletion^39^. Our previous study demonstrated that AAV-PERM1 enhances mitochondrial biogenesis in healthy C57BL/6 mouse hearts, as evidenced by increased mtDNA copy number and upregulation of PGC-1α^22^. To investigate whether AAV-PERM1 preserves mitochondrial biogenesis under pressure overload, we quantified mtDNA copy number in Sham and TAC hearts from AAV-GFP and AAV-PERM1-treated mice. TAC led to a ~50% reduction in mtDNA (normalized to nuclear DNA) in GFP hearts (p<0.05, **Fig. 3P**). AAV-PERM1 significantly increased mtDNA copy number compared to GFP Sham (1.0 vs. 1.5, p<0.05) and maintained it during TAC (1.5 in Sham vs. 1.3 in TAC, p>0.05). This was consistent with Western blot data showing that TFAM, a key transcription factor for mtDNA replication, was downregulated by TAC in GFP hearts but upregulated by AAV-PERM1 in both Sham and TAC conditions (136% and 174% of GFP Sham, respectively, p<0.05, **Fig. 3A-B**).

To assess whether AAV-PERM1 influences mitophagy as a mechanism for preserving mitochondrial content, we examined protein levels of OPA1 and mitofusin 1 (MFN1)—key regulators of mitochondrial fusion and fission. TAC did not alter the expression of these proteins in GFP-treated hearts (p>0.05), but AAV-PERM1 significantly increased OPA1 expression in both Sham and TAC groups (**Fig. 3Q-S**). Similarly, neither TAC nor AAV-PERM1 affected the total expression levels of dynamin-related protein 1 (DRP1), a key GTPase that mediates mitochondrial fission (**Fig. 3Q,T**). Notably, DRP1 phosphorylation was elevated by TAC in both GFP and AAV-PERM1 groups to a similar extent (**Fig. 3Q, U**), indicating DRP1 activation occurred in response to pressure overload and was not further enhanced by AAV-PERM1.

In summary, these results, together with data presented in **Fig. 2**, demonstrate that AAV-PERM1 preserves both mitochondrial biogenesis and bioenergetics in the pressure-overloaded heart. This protective effect is associated with the maintenance or upregulation of key transcriptional regulators and structural components of mitochondrial oxidative phosphorylation, while AAV-PERM1 had little effect on mitophagy.

### Metabolomics analysis reveals a distinct profile in AAV-PERM1 hearts during pressure overload

To determine whether preserved mitochondrial bioenergetics by AAV-PERM1 under pressure overload altered the cardiac metabolome, we performed untargeted metabolomics on cardiac tissue from AAV-GFP and AAV-PERM1 mice subjected to Sham or TAC surgery (8 weeks post-surgery). Principal component analysis (PCA) revealed a subtle but statistically significant difference between GFP Sham and GFP TAC hearts (p<0.05, **Fig. 4A**), indicating that TAC induces global changes in the cardiac metabolomic profile. Integrated pathway enrichment and topology analysis using MetaboAnalyst 6.0 identified several metabolic pathways significantly affected by TAC, including ketone body metabolism, the tricarboxylic acid (TCA) cycle, carnitine synthesis, and fatty acid and pyruvate metabolism (**Fig. 4B-C**). In contrast, the PCA plot comparing PERM1 Sham and PERM1 TAC hearts showed no significant separation (p<0.05, **Fig. 4D**), although modest changes were observed in linoleic acid metabolism and amino acid metabolism (**Fig. 4D-E**). Furthermore, the 3D PCA plot comparing GFP TAC and PERM1 TAC hearts revealed distinct metabolic profiles (**Fig. 4F**). Specifically, TAC-induced accumulation of spermidine and TCA cycle intermediates, including succinate, fumarate, and malate, was normalized by AAV-PERM1 (**Fig. 4G-H**). Collectively, these metabolomic analyses demonstrate that TAC disrupts multiple metabolic pathways converging on oxidative phosphorylation (OXPHOS), and that AAV-PERM1 partially restores metabolic homeostasis under pressure overload conditions.

**Figure 4.**
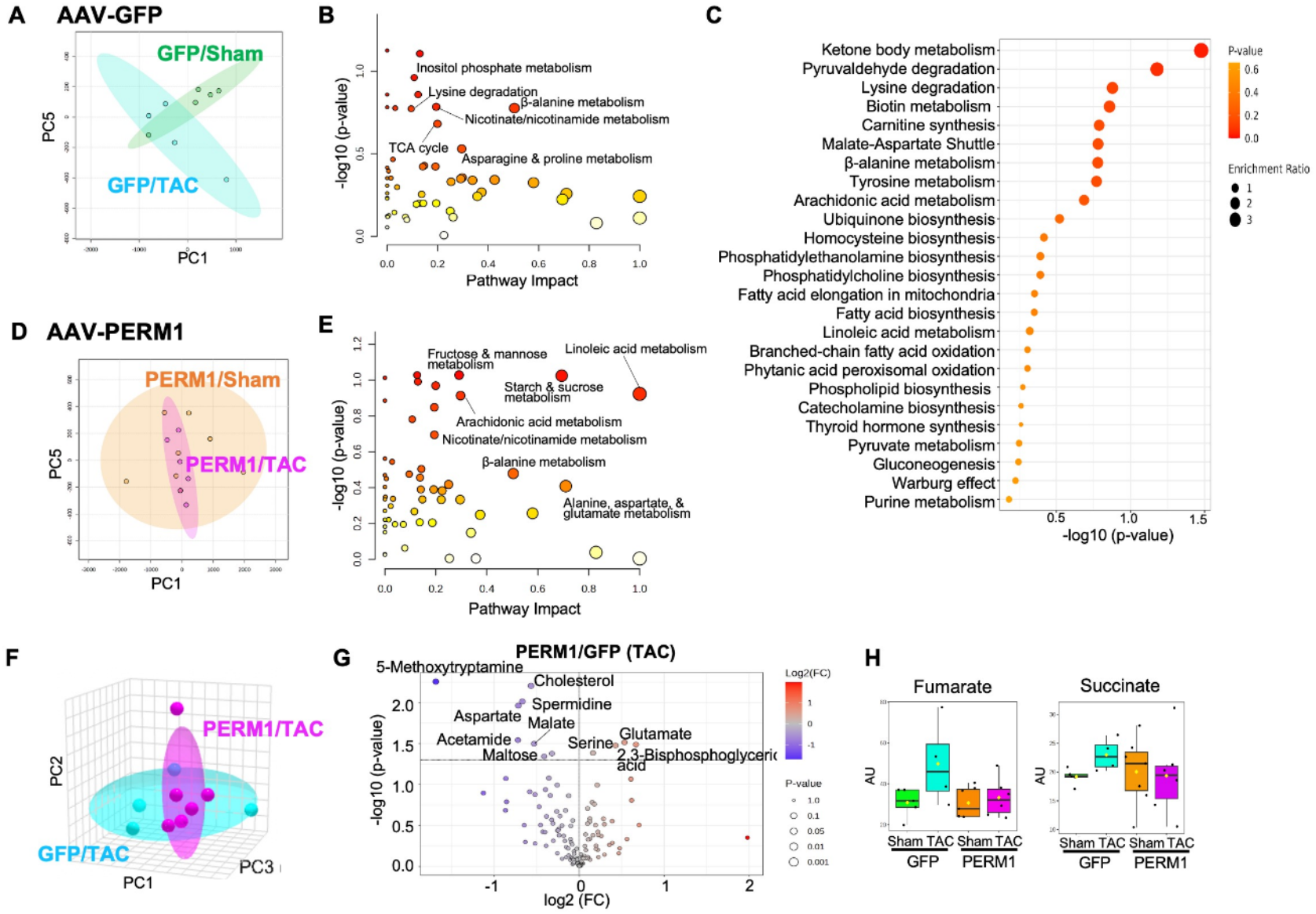
Metabolomic profiling of AAV-GFP and AAV-PERM1 hearts subjected to Sham or TAC. **A– C**, Metabolomic profiles of AAV-GFP Sham and AAV-GFP TAC hearts. The principal component analysis (PCA) plot shows a distinct separation between TAC and Sham groups, indicating altered metabolomic profiles in response to TAC (**A**, p<0.05). Pathway analysis (**B**) and enrichment analysis (**C**) highlight specific metabolic pathways affected by TAC. **D–E**, Comparison of AAV-PERM1 Sham and AAV-PERM1 TAC hearts. The PCA plot shows no significant separation between the two groups (**D**, p>0.05), suggesting that AAV-PERM1 preserves a metabolomic profile in TAC hearts comparable to Sham. **F–H**, Comparison of AAV-GFP TAC and AAV-PERM1 TAC hearts. A distinct separation in metabolomic profiles is observed (**F**). TAC-induced accumulation of TCA cycle intermediates (malate, fumarate, succinate) and spermidine is attenuated in AAV-PERM1 hearts (**G–H**).

### AAV-PERM1 prevents excessive O-GlcNAcylation in the heart during pressure overload

O-GlcNAcylation is a post-translational modification of proteins that is regulated by two enzymes: O-GlcNAc transferase (OGT) that adds O-GlcNAc to proteins; O-GlcNAcase (OGA) that removes O-GlcNAc from proteins. Excessive or prolonged activation of O-GlcNAcylation is associated with heart failure and hypertrophy^40–50^. We previously demonstrated that PERM1 negatively regulates O-GlcNAcylation by repressing *Ogt* expression through transcriptional regulation, and enhances FAO by preventing O-GlcNAcylation of PGC-1α, thus preserving its interaction with PPARα, a key transcription factor of FAO^34^. To determine whether the cardioprotective effect and enhanced mitochondrial respiration observed with AAV-PERM1 (**Fig. 1–2**) involves suppression of excessive O-GlcNAcylation, we performed western blot analysis to assess total O-GlcNAcylated proteins and the expression levels of OGT and OGA. In agreement with previous studies in human failing hearts and TAC mouse models, 8 weeks of TAC induced a significant increase in both high molecular weight (HMW) and low molecular weight (LMW) fractions of total O-GlcNAcylated proteins (**Fig. 5A-C)** and upregulation of OGT (**Fig. 5D**). AAV-PERM1 treatment prior to TAC prevented the increase in both O-GlcNAcylation and OGT expression, maintaining levels comparable to baseline (**Fig. 5**). There was a trend towards increased OGA expression in AAV-PERM1 Sham mice, which did not reach statistical significance due to a large dispersion in outcomes, but otherwise the OGA expression was minimally affected by either TAC or AAV-PERM1 (**Fig. 5E**). Importantly, O-GlcNAcylation of PGC-1α was markedly increased following TAC and was completely suppressed below basal levels by AAV-PERM1 (**Fig. 5F-G**). Given that transgenic mice overexpressing OGA has been shown to confer resistant to pressure overload by reducing O-GlcNAcylation and restoring Complex I activity^49^, these data suggest that AAV-PERM1 protects against mitochondrial impairment and cardiac dysfunction, at least in part, by preventing pathological increases in O-GlcNAcylation during pressure overload.

**Figure 5.**
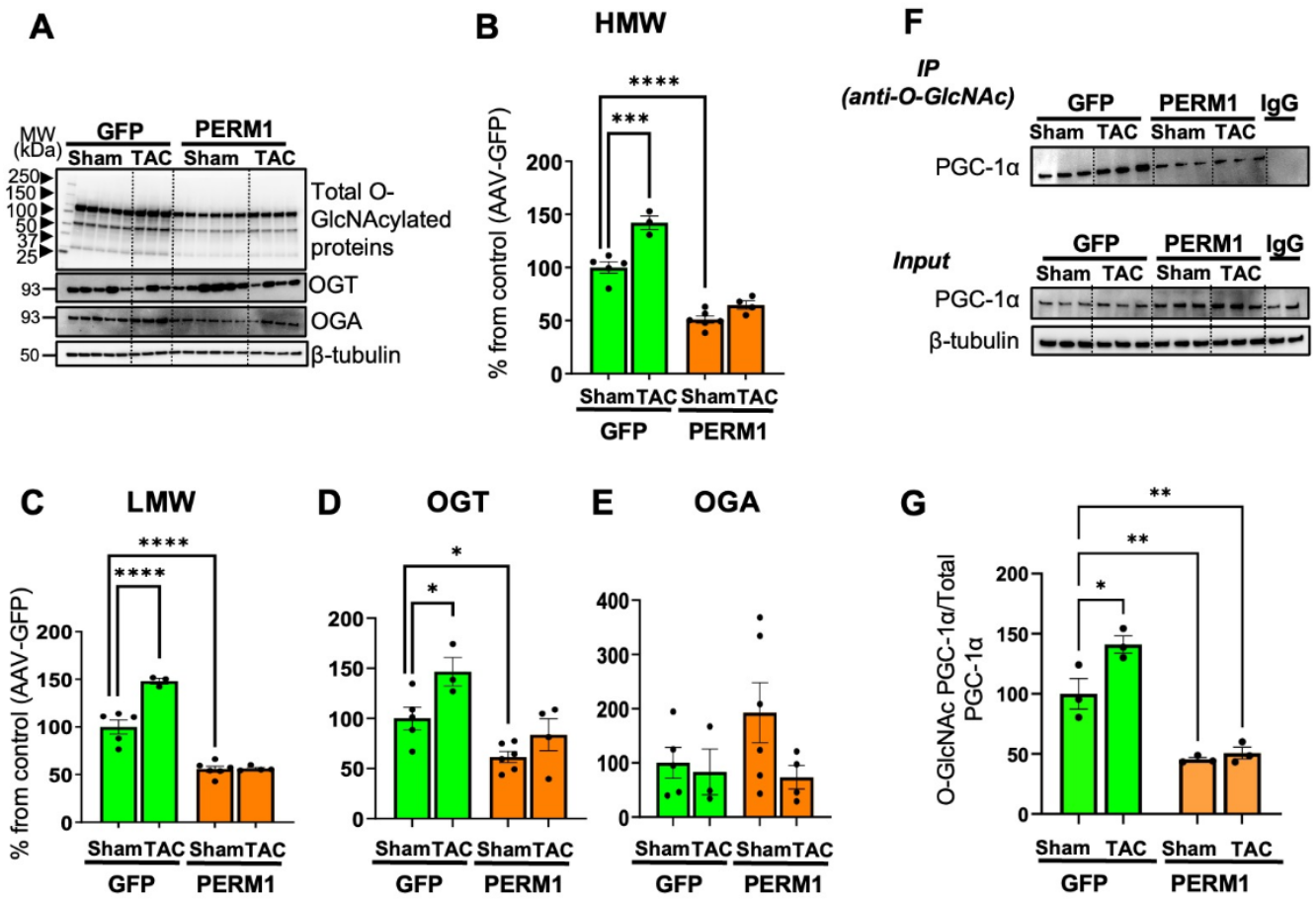
AAV-PERM1 prevents TAC-induced elevation of O-GlcNAcylation. Western blot analysis shows a significant increase in total O-GlcNAcylated proteins at both high molecular weight (HMW, **Panel B**) and low molecular weight (LMW, **Panel C**) ranges, as well as increased OGT expression in GFP-TAC hearts compared to GFP-Sham hearts (**D**), with no significant change in OGA levels (**E**). In contrast, AAV-PERM1 reduced total O-GlcNAcylated proteins of both HMW and LMW (**B-C**) and OGT expression (**D**), and these suppressed levels were maintained during pressure overload. O-GlcNAcylated proteins were quantified separately for HMW (≥75 kDa) and LMW (<75 kDa) fractions (see Methods for details). **F-G**, Statistical comparisons presented are results of post-hoc pairwise Holm-Sidak test applied after two-way ANOVA. ****: p<0.0001, ***: p<0.001, *: p<0.05, mean ± SEM

### PERM1 interacts with creatine kinase and Troponin C

Our previous study demonstrated that AAV-mediated overexpression of PERM1 enhances cardiac contractility in healthy hearts that are not limited by ATP availability [22], suggesting that PERM1 may function beyond simply promoting ATP production. Creatine kinase (CK) plays a key role in shuttling ATP produced in mitochondria to sites of ATP utilization. In the heart, approximately 90% of mitochondrial ATP is consumed to support contractility ^51^. We hypothesized that PERM1 not only enhances mitochondrial oxidative phosphorylation but also improves the efficiency of coupling between ATP generation and its utilization in the sarcomere. To test whether PERM1 interacts with CK in the heart, we performed co-immunoprecipitation (Co-IP) assays using WT heart tissue. Pulldown with anti-CK antibodies revealed that CK interacts with PERM1 (**Fig. 6A**). Furthermore, pulldown assays using both anti-CK and anti-PERM1 antibodies demonstrated that troponin C (TnC), a key component of the contractile apparatus in the sarcomere, forms complexes with both CK and PERM1 in WT and AAV-PERM1-treated hearts (**Fig. 6B**). In contract, neither the total expression levels and activity of CK was changed by loss- and gain-of-function of PERM1 in mice (**Fig. S1**). These results suggest that PERM1 promotes the spatial coupling of CK with TnC, potentially enhancing the efficiency of ATP delivery to the sarcomere and thereby supporting contractile function.

**Figure 6.**
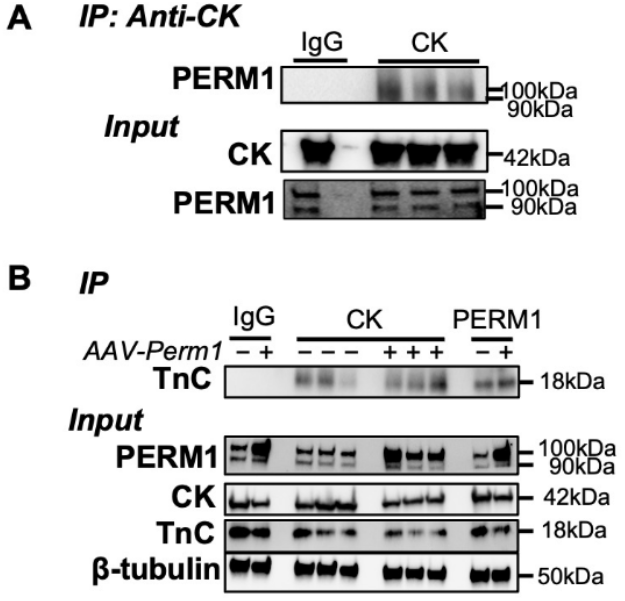
PERM1 interacts with creatine kinase (CK) and Troponin C (TnC). **A-B**, Co-immunoprecipitation (Co-IP) assays showing that PERM1 interacts with CK in WT hearts (**A**), and that TnC forms complexes with both CK and PERM1 in WT and AAV-PERM1-treated hearts (**B**).

### Cardiac-Specific AAV-PERM1 (TNNT2) Improves Heart Function and Mitochondrial Biogenesis with limited Gene Delivery Efficiency

The prominent cardioprotective effects shown in Fig. 1–4 were obtained using our first-generation AAV-PERM1 vector containing the cytomegalovirus (CMV) promoter, a strong, constitutive viral promoter widely used in AAV vectors to drive high levels of transgene expression across various cell types. While we confirmed that this CMV-driven vector achieves robust PERM1 overexpression in mouse hearts without altering expression in skeletal muscle or adipose tissue^22^, cardiac-specific gene delivery is critical for translational applications. To address this, we developed a second-generation AAV-PERM1 vector using the TNNT2 promoter, which is active exclusively in cardiac tissue (AAV-PERM1/TNNT2). Consistent with our previous study^22^ and current results (**Fig. 1**), echocardiography demonstrated that a 4-week treatment with AAV-PERM1/TNNT2 significantly increased LVEF in the heart compared to baseline (**Fig. S2A**); however, the LVEF increase was stronger with the CMV promoter (**Fig. S2B**). Similarly, mitochondrial DNA (mtDNA) copy number was significantly increased by AAV-PERM1/TNNT2, but its capacity to enhance mitochondrial biogenesis was lower than that of AAV-PERM1/CMV (1.5-fold vs. 3.8-fold increase over GFP, **Fig. S2C**). Western blot analysis further confirmed that AAV-PERM1/TNNT2 increased PERM1 protein levels in the heart by approximately 1.3-fold, while AAV-PERM1/CMV resulted in a much greater increase (8-fold, **Fig. S2D–E**). Importantly, neither AAV-PERM1/TNNT2 nor AAV-PERM1/CMV altered PERM1 expression in skeletal muscle (data not shown), confirming cardiac-specific overexpression in both constructs.

## 4. Discussion

In this study, we demonstrate that AAV-mediated gene delivery of PERM1 (AAV-PERM1) fully prevents the onset of heart failure in response to pressure overload by cardiac contractility, mitochondrial biogenesis, and mitochondrial respiratory capacity. The cardioprotective effects of AAV-PERM1 were also associated with suppression of cardiac hypertrophy and fibrosis. Furthermore, we show that PERM1 interacts with creatine kinase (CK), potentially facilitating the coupling between ATP production and contractile function.

AAV vectors have emerged as powerful tools for gene therapy due to their ability to provide sustained gene expression with low immunogenicity. Over the past decade, AAV-mediated cardiac gene therapy has shifted the paradigm from symptomatic treatment to disease-modifying—and potentially curative— strategies by targeting molecular drivers of cardiac remodeling. Despite this promise, no FDA-approved AAV therapies for heart failure currently exist, partly due to the lack of validated, disease-modifying gene targets. PERM1, a striated muscle-specific protein induced by PGC-1α and ERRs, is highly expressed in the heart ^16^. We previously identified its downregulation in both human HFrEF and the TAC mouse model^15^. Unlike other essential regulators of mitochondrial bioenergetics, PERM1 deletion is not embryonically lethal; Perm1-KO mice exhibit only mild systolic dysfunction and reduced mitochondrial energetics ^16^. These characteristics make PERM1 an attractive therapeutic target for fine-tuning mitochondrial function without disrupting physiological homeostasis. To evaluate its therapeutic potential, we generated AAV-PERM1, which achieved ~3-fold overexpression of PERM1 in adult C57BL/6 mouse hearts ^22^. In healthy hearts, this overexpression enhanced systolic function and mitochondrial biogenesis without any evidence of adverse myocardial remodeling or any other side effects ^22^. These findings are particularly notable given that PGC-1α overexpression in transgenic mice leads to uncontrolled mitochondrial proliferation and cardiomyopathy^10^. In the current study, AAV-PERM1 expression remained stable during pressure overload which fully prevented the development of HFrEF. These effects were consistent with—but exceeded—those observed in transgenic mice with constitutive PERM1 overexpression ^52^, confirming the therapeutic potential of PERM1 and establishing AAV-PERM1 as a practical and translationally relevant gene therapy strategy.

PERM1 has been identified as a regulator of mitochondrial bioenergetics. In vitro, we found that PERM1 knockdown in cardiomyocytes and H9c2 cells reduces oxidative capacity with both pyruvate and palmitate substrates, whereas adenoviral overexpression of PERM1 enhances oxidative capacity for both ^34^. However, whether PERM1 can increase mitochondrial respiratory capacity *in vivo* remained unknown. In addition, although transgenic (Tg) mice overexpressing PERM1 were protected against systolic dysfunction during pressure overload, most components of the electron transport chain (ETC) and the MICOS-MIB complex were unaffected by TAC in this model ^52^. Thus, it was unclear whether PERM1 overexpression could prevent mitochondrial impairment in pressure-overloaded hearts. To address these questions, we employed a comprehensive approach combining Oroboros O2k high-resolution respirometry with the Horiba QuantMaster to simultaneously assess O_2_ consumption, H_2_O_2_ production, and electron leak in mitochondria isolated from the same hearts. Our results demonstrate that AAV-PERM1 enhances mitochondrial respiration supported both pyruvate/malate and octanoylcarnitine/malate substrates. This is the first study to show that AAV-PERM1 improves mitochondrial function in the intact heart. Notably, enhanced pyruvate-driven respiration was preserved even under TAC, whereas fatty acid–driven respiration returned to baseline. These findings suggest that PERM1 exerts substrate-specific effects, preferentially supporting TCA cycle metabolism under stress. Enhanced mitochondrial conductance under both conditions further supports a role for PERM1 in maintaining mitochondrial quality.

Analysis of H_2_O_2_ emission (JH2O2) and electron leak revealed that PERM1 protects against oxidative stress predominantly in the pyruvate/malate condition. In contrast, fatty acid-driven respiration under TAC resulted in elevated electron leak at high respiratory demand, suggesting a maladaptive response. These findings support the idea that PERM1 enhances metabolic resilience in the failing heart by maintaining redox balance and mitochondrial integrity, especially through glucose/TCA metabolism. Further research is needed to delineate the underlying mechanisms, particularly given that electron transport chain function declines with age and in diabetic cardiomyopathy ^53^.

Our studies also show that AAV-PERM1 promotes mitochondrial biogenesis, evidenced by a strong association between PERM1 expression and mtDNA content ^22^ (**Fig. 2**). This is particularly relevant since effective HFrEF therapies must address mitochondrial dysfunction, which is often independent of PGC-1α. In both our model and in human HFrEF, mtDNA-encoded OXPHOS gene expression is reduced despite unchanged PGC-1α levels ^39^. Although AAV-PERM1 modestly upregulated PGC-1α in both control and TAC hearts, it is unlikely that this contributed significantly to the observed protection, especially since moderate PGC-1αα overexpression in TAC mice fails to preserve systolic function or respiration ^54^.

The mechanisms by which PERM1 regulates mitochondrial energetics are only beginning to be understood. We previously demonstrated that PERM1 acts as a transcriptional coactivator in the nucleus, interacting with ERRα and PGC-1α to drive the expression of mitochondrial genes [15]. Other studies have reported that PERM1 promotes fatty acid oxidation (FAO) through interaction with PPARα. In the current study, we show that AAV-PERM1 upregulates ERRα and PPARα *in vivo*, consistent with enhanced mitochondrial function. Supporting these findings, our western blot data demonstrate that key transcription factors regulating oxidative phosphorylation (OXPHOS) genes (ERRα, NRF1) and FAO (PPARαα) were downregulated in GFP-TAC hearts but were maintained at baseline levels or above in AAV-PERM1-treated hearts (**Fig. 3**). Similarly, cytochrome c (CytC) and ATP synthase β, both components of the electron transport chain, showed parallel changes in response to TAC and AAV-PERM1 (**Fig. 3**). These results suggest that AAV-PERM1 enhances mitochondrial energetics by normalizing OXPHOS gene expression during pressure overload through nuclear receptor pathways. Unexpectedly, expression levels of key FAO enzymes (CPT1b, CPT2, MCAD) remained unchanged in response to either TAC or AAV-PERM1 treatment (**Fig. 3**). Given the observed reduction in octanoylcarnitine (Oct/M)-stimulated respiration in TAC hearts and its rescue by AAV-PERM1 (**Fig. 2**), it is possible that the FAO bottleneck lies in the altered expression and/or function of mitochondrial trifunctional protein (MTP), a complex of three enzymes involved in β-oxidation within the mitochondrial matrix. Future RNA-seq and proteomics analyses will be required to identify the precise molecular bottlenecks in FAO in TAC hearts and the regulatory targets normalized by AAV-PERM1 during pressure overload.

We also previously demonstrated that PERM1 suppresses O-GlcNAcylation—possibly by reducing GLUT1/4 expression and limiting glucose entry into the hexosamine biosynthesis pathway ^34^. Loss of PERM1 results in O-GlcNAcylation of PGC-1α, disrupting its interaction with PPARα and impairing FAO. Here, AAV-PERM1 prevented TAC-induced O-GlcNAcylation, suggesting that preserved PGC-1α /PPARα signaling contributes to its cardioprotective effects. In addition to its nuclear function, PERM1 localizes to the mitochondrial outer membrane, where it may stabilize mitochondria–sarcolemma coupling via ankyrin B and the MICOS-MIB complex ^55^. Whether this contributes to mitochondrial stability during pressure overload remains to be investigated.

Our gain- and loss-of-function studies also support a role for PERM1 in contractility. AAV-PERM1 achieving ~3-fold overexpression of PERM1, enhances cardiac contractility in healthy hearts which are not ATP-limited, suggesting a role beyond energy production. Here, we show that PERM1 interacts with CK and localizes to the sarcomere, likely facilitating energy transfer and coupling metabolism to mechanical function. This points to a novel role for PERM1 in cardiac mechano-energetics. Future studies using PERM1 mutants lacking the nuclear localization domain will help distinguish the contribution of nuclear vs. cytoplasmic functions. Moreover, previous studies in C2C12 cells showed that PERM1 regulates CaMKIIβ via the MAPK pathway. Whether PERM1 modulates calcium handling or CaMKII activation in cardiomyocytes during stress remains to be determined.

Importantly, AAV-PERM1 also attenuates TAC-induced cardiac fibrosis and hypertrophy. While PERM1’s direct role in fibrotic signaling is unclear, our RNA-seq data show that PERM1 knockdown upregulates pro-fibrotic genes such as Col3a1 and Col5a2—both induced by TAC and suppressed by AAV-PERM1. Although PERM1 is expressed in neonatal cardiac fibroblasts (unpublished), its role in adult fibroblasts remains unknown. Regarding hypertrophy, we previously reported that PERM1 overexpression downregulates GLUT4, with limited enhancement of glucose-driven respiration compared to pyruvate. Given that the pentose phosphate pathway (PPP)—an accessory glucose pathway—is often upregulated in hypertrophy and supports nucleotide synthesis, it is plausible that AAV-PERM1 limits hypertrophy by restricting glucose uptake and PPP activation, although this remains to be confirmed.

In conclusion, this study identifies PERM1 as a promising therapeutic target for HFrEF. AAV-PERM1 gene therapy preserves mitochondrial biogenesis, enhances mitochondrial function, improves contractility, and reduces fibrosis—without inducing adverse remodeling. These findings establish a strong foundation for the development of PERM1-based gene therapies aimed at restoring energetic and mechanical homeostasis in the failing heart.

## Data availability

All measured data points necessary to support the conclusions of this article are included in the manuscript. Raw data (echocardiography images, original blots, original GC-MS runs files, and others) are available from authors upon a reasonable request.

## Funding information

This study was supported by NIH R01HL156667 (J.S.W.), FBRI Virginia Tech Carilion Operation Fund (J.S.W.), Seale Innovation Fund (J.S.W.), American Heart Association (AHA) Postdoctoral Fellowship 24POST1186520 (K.S.), Virginia Tech Research and Innovation Postdoctoral Scholarship (K.S.), NIH R01AR050429 (Z.Y.), NIH R01AR077440 (Z.Y.) and a grant by Red Gates Foundation (Z.Y.).

## Competing Interest Statement

J.S.W, A.V.Z, and K.S. are co-founders of a startup company **Vectorial Corp**. created for the purpose of translation of AAV-PERM1 vector therapy into clinical practice.

## Declaration of Generative AI and AI-assisted technologies in the writing process

The authors did not use generative AI or AI-assisted technologies in the development of this manuscript.

## References

1. Heidenreich, P.A., et al. 2022 AHA/ACC/HFSA Guideline for the Management of Heart Failure: A Report of the American College of Cardiology/American Heart Association Joint Committee on Clinical Practice Guidelines. Circulation 145, e895–e1032 (2022).

2. Ghionzoli, N., et al. Current and emerging drug targets in heart failure treatment. Heart Fail Rev 27, 1119–1136 (2022).

3. Ingwall, J.S. & Weiss, R.G. Is the failing heart energy starved? On using chemical energy to support cardiac function. Circ Res 95, 135–145 (2004).

4. Arany, Z., et al. Transverse aortic constriction leads to accelerated heart failure in mice lacking PPARα-gamma coactivator 1alpha. Proc Natl Acad Sci U S A 103, 10086–10091 (2006).

5. Huss, J.M., et al. The nuclear receptor ERRαalpha is required for the bioenergetic and functional adaptation to cardiac pressure overload. Cell Metab 6, 25–37 (2007).

6. Oka, S., et al. PPARαalpha-Sirt1 complex mediates cardiac hypertrophy and failure through suppression of the ERRα transcriptional pathway. Cell Metab 14, 598–611 (2011).

7. Huss, J.M. & Kelly, D.P. Nuclear receptor signaling and cardiac energetics. Circ Res 95, 568–578 (2004).

8. Rowe, G.C., Jiang, A. & Arany, Z. PGC-1α coactivators in cardiac development and disease. Circulation research 107, 825–838 (2010).

9. Kolwicz, S.C., Jr. & Tian, R. Glucose metabolism and cardiac hypertrophy. Cardiovasc Res 90, 194–201 (2011).

10. Lehman, J.J., et al. Peroxisome proliferator-activated receptor gamma coactivator-1 promotes cardiac mitochondrial biogenesis. J Clin Invest 106, 847–856 (2000).

11. Karamanlidis, G., Garcia-Menendez, L., Kolwicz, S.C., Jr., Lee, C.F. & Tian, R. Promoting PGC1alpha-driven mitochondrial biogenesis is detrimental in pressure-overloaded mouse hearts. Am J Physiol Heart Circ Physiol 307, H1307–1316 (2014).

12. Young, M.E., Laws, F.A., Goodwin, G.W. & Taegtmeyer, H. Reactivation of peroxisome proliferator-activated receptor alpha is associated with contractile dysfunction in hypertrophied rat heart. J Biol Chem 276, 44390–44395 (2001).

13. Lasheras, J., et al. Cardiac-Specific Overexpression of ERRαγ in Mice Induces Severe Heart Dysfunction and Early Lethality. Int J Mol Sci 22(2021).

14. Cho, Y., Hazen, B.C., Russell, A.P. & Kralli, A. Peroxisome proliferator-activated receptor gamma coactivator 1 (PGC-1α)- and estrogen-related receptor (ERRα)-induced regulator in muscle 1 (Perm1) is a tissue-specific regulator of oxidative capacity in skeletal muscle cells. J Biol Chem 288, 25207–25218 (2013).

15. Oka, S.I., et al. Perm1 regulates cardiac energetics as a downstream target of the histone methyltransferase Smyd1. PLoS One 15, e0234913 (2020).

16. Oka, S.I., et al. PERM1 regulates energy metabolism in the heart via ERRαalpha/PGC-1αalpha axis. Front Cardiovasc Med 9, 1033457 (2022).

17. Yamada, K.P., Tharakan, S. & Ishikawa, K. Consideration of clinical translation of cardiac AAV gene therapy. Cell Gene Ther Insights 6, 609–615 (2020).

18. Sahoo, S., Kariya, T. & Ishikawa, K. Targeted delivery of therapeutic agents to the heart. Nat Rev Cardiol 18, 389–399 (2021).

19. Mazurek, R., et al. AAV delivery strategy with mechanical support for safe and efficacious cardiac gene transfer in swine. Nat Commun 15, 10450 (2024).

20. Ishikawa, K., et al. Cardiac I-1c overexpression with reengineered AAV improves cardiac function in swine ischemic heart failure. Mol Ther 22, 2038–2045 (2014).

21. Hammoudi, N., Ishikawa, K. & Hajjar, R.J. Adeno-associated virus-mediated gene therapy in cardiovascular disease. Curr Opin Cardiol 30, 228–234 (2015).

22. Sreedevi, K., et al. Adeno-associated virus-mediated gene delivery of Perm1 enhances cardiac contractility in mice. Am J Physiol Heart Circ Physiol 327, H1112–h1118 (2024).

23. Riehle, C., et al. PGC-1αbeta deficiency accelerates the transition to heart failure in pressure overload hypertrophy. Circ Res 109, 783–793 (2011).

24. Drake, J.C., et al. Mitochondria-localized AMPK responds to local energetics and contributes to exercise and energetic stress-induced mitophagy. Proc Natl Acad Sci U S A 118(2021).

25. Fisher-Wellman, K.H., et al. Mitochondrial Diagnostics: A Multiplexed Assay Platform for Comprehensive Assessment of Mitochondrial Energy Fluxes. Cell Rep 24, 3593–3606 e3510 (2018).

26. Tarpey, M.D., Amorese, A.J., Balestrieri, N.P., Fisher-Wellman, K.H. & Spangenburg, E.E. Doxorubicin causes lesions in the electron transport system of skeletal muscle mitochondria that are associated with a loss of contractile function. J Biol Chem 294, 19709–19722 (2019).

27. Fisher-Wellman, K.H., et al. Mitochondrial glutathione depletion reveals a novel role for the pyruvate dehydrogenase complex as a key H2O2-emitting source under conditions of nutrient overload. Free Radic Biol Med 65, 1201–1208 (2013).

28. Fisher-Wellman, K.H., et al. Pyruvate dehydrogenase complex and nicotinamide nucleotide transhydrogenase constitute an energy-consuming redox circuit. Biochem J 467, 271–280 (2015).

29. Warren, J.S., et al. Histone methyltransferase Smyd1 regulates mitochondrial energetics in the heart. Proc Natl Acad Sci U S A (2018).

30. Shibayama, J., et al. Metabolic determinants of electrical failure in ex-vivo canine model of cardiac arrest: evidence for the protective role of inorganic pyrophosphate. PLoS One 8, e57821 (2013).

31. Shibayama, J., et al. Metabolic remodeling in moderate synchronous versus dyssynchronous pacing-induced heart failure: integrated metabolomics and proteomics study. PLoS One 10, e0118974 (2015).

32. Richards, D.A., et al. Distinct Phenotypes Induced by Three Degrees of Transverse Aortic Constriction in Mice. Sci Rep 9, 5844 (2019).

33. Wang, X., et al. A time-series minimally invasive transverse aortic constriction mouse model for pressure overload-induced cardiac remodeling and heart failure. Front Cardiovasc Med 10, 1110032 (2023).

34. Sreedevi, K., et al. PERM1 regulates mitochondrial energetics through O-GlcNAcylation in the heart. J Mol Cell Cardiol 198, 1–12 (2025).

35. Bugger, H., et al. Proteomic remodelling of mitochondrial oxidative pathways in pressure overload-induced heart failure. Cardiovasc Res 85, 376–384 (2010).

36. Tsutsui, H., Kinugawa, S. & Matsushima, S. Oxidative stress and heart failure. Am J Physiol Heart Circ Physiol 301, H2181–2190 (2011).

37. Villena, J.A. New insights into PGC-1α coactivators: redefining their role in the regulation of mitochondrial function and beyond. FEBS J 282, 647–672 (2015).

38. Finck, B.N. & Kelly, D.P. PGC-1α coactivators: inducible regulators of energy metabolism in health and disease. J Clin Invest 116, 615–622 (2006).

39. Karamanlidis, G., et al. Defective DNA replication impairs mitochondrial biogenesis in human failing hearts. Circ Res 106, 1541–1548 (2010).

40. Clark, R.J., et al. Diabetes and the accompanying hyperglycemia impairs cardiomyocyte calcium cycling through increased nuclear-GlcNAcylation. Journal of Biological Chemistry 278, 44230–44237 (2003).

41. Erickson, J.R., et al. Diabetic hyperglycaemia activates CaMKII and arrhythmias by O-linked glycosylation. Nature 502, 372–376 (2013).

42. Jennifer L. McLarty, S.A.M., and John C. Chatham. Post-translational protein modification by O-linked N-acetylglucosamine: Its role in mediating the adverse effects of diabetes on the heart. J Biol Chem (2013).

43. Marsh, S.A., Dell’Italia, L.J. & Chatham, J.C. Activation of the hexosamine biosynthesis pathway and protein O-GlcNAcylation modulate hypertrophic and cell signaling pathways in cardiomyocytes from diabetic mice. Amino Acids 40, 819–828 (2011).

44. Lunde, I.G., et al. Cardiac O-GlcNAc signaling is increased in hypertrophy and heart failure. Physiol Genomics 44, 162–172 (2012).

45. Ma, J. & Hart, G.W. Protein O-GlcNAcylation in diabetes and diabetic complications. Expert Rev Proteomics 10, 365–380 (2013).

46. Facundo, H.T., et al. O-GlcNAc signaling is essential for NFAT-mediated transcriptional reprogramming during cardiomyocyte hypertrophy. Am J Physiol-Heart C 302, H2122–H2130 (2012).

47. Gélinas, R., et al. AMPK activation counteracts cardiac hypertrophy by reducing O-GlcNAcylation. Nature Communications 9(2018).

48. Chen, X.L., et al. Increased O-GlcNAcylation induces myocardial hypertrophy. In Vitro Cell Dev-An 56, 735–743 (2020).

49. Umapathi, P., et al. Excessive O-GlcNAcylation Causes Heart Failure and Sudden Death. Circulation 143, 1687–1703 (2021).

50. Matsuno, M., et al. -GlcNAcylation-induced GSK-3β activation deteriorates pressure overload-induced heart failure lack of compensatory cardiac hypertrophy in mice. Front Endocrinol 14(2023).

51. Khan, M.S. & Butler, J. Targeting Mitochondrial Function in Heart Failure: Makes Sense But Will it Work? JACC Basic Transl Sci 4, 158–160 (2019).

52. Tachibana, S., et al. Perm1 Protects the Heart From Pressure Overload-Induced Dysfunction by Promoting Oxidative Metabolism. Circulation 147, 916–919 (2023).

53. Lesnefsky, E.J., Chen, Q. & Hoppel, C.L. Mitochondrial Metabolism in Aging Heart. Circ Res 118, 1593–1611 (2016).

54. Pereira, R.O., et al. Maintaining PGC-1αalpha expression following pressure overload-induced cardiac hypertrophy preserves angiogenesis but not contractile or mitochondrial function. FASEB J 28, 3691–3702 (2014).

55. Bock, T., et al. PERM1 interacts with the MICOS-MIB complex to connect the mitochondria and sarcolemma via ankyrin B. Nat Commun 12, 4900 (2021).

