## Supplemental Figures for "PERM1 Gene Delivery via AAV Prevents Heart Failure in a Mouse Model of Pressure Overload"

### SUPPLEMENTARY FIGURES.

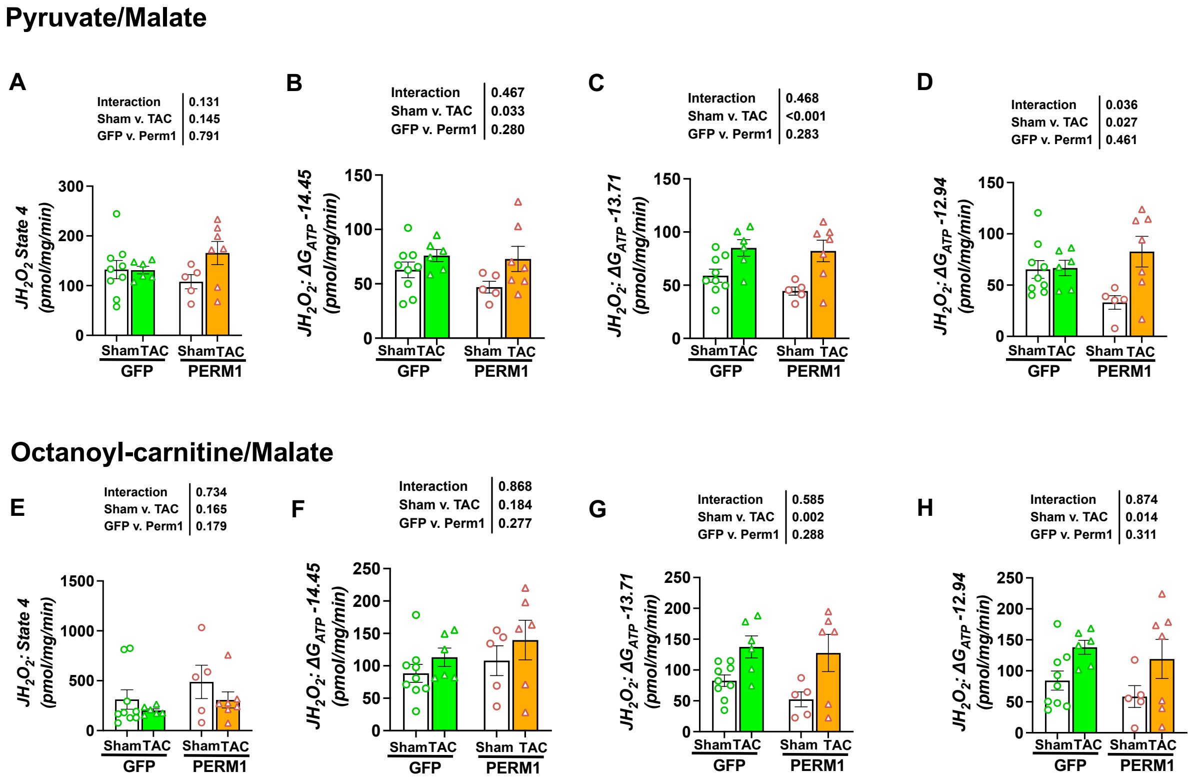


**Figure S1**. **H2O2 flux (JH2O2) as a function of substrate and energy demand.** (**A-D**), JH2O2 in State 4 and at different levels of $\Delta G_{ATP}$ under pyruvate/malate conditions. (**E-H**), JH2O2 in State 4 and at different levels of $\Delta G_{ATP}$ under octanoyl-carnitine/malate conditions. Under TAC, there is an overall trend of JH2O2 increase with increasing energy demand, which is not affected by PERM1 overexpression.

### Pyruvate/Malate

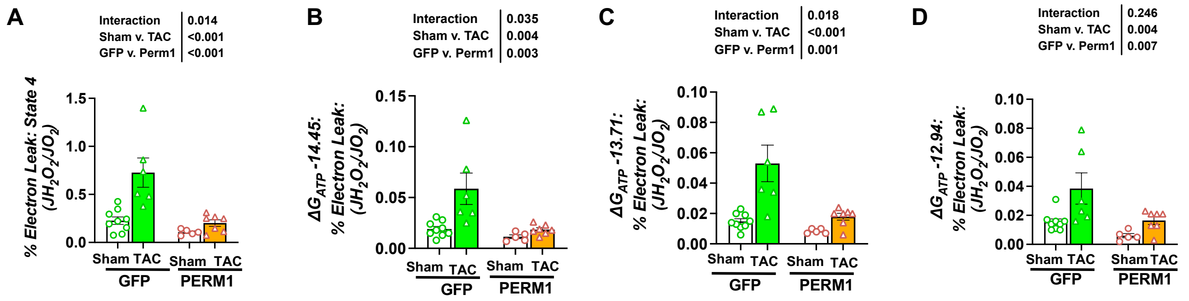


### Octanoyl-carnitine/Malate

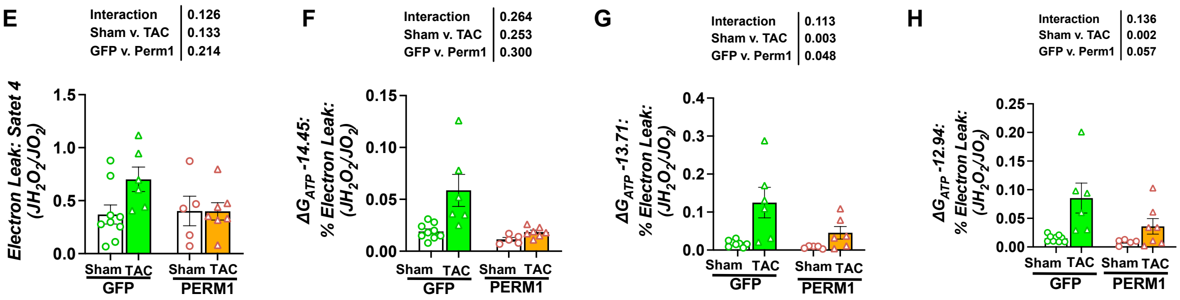


**Figure S2. Percent electron leak (JH2O2/JO2) as a function of substrate and energy demand.** (**A-D**), % electron leak in State 4 and at different levels of $\Delta G_{ATP}$ under pyruvate/malate conditions. (**E-H**), % electron leak in State 4 and at different levels of $\Delta G_{ATP}$ under octanoyl-carnitine/malate conditions. Pressure overload significantly increases % electron leak under all tested conditions. This increase is fully or partially prevented by PERM1 overexpression with a trend of less protection at higher levels of energy demand.

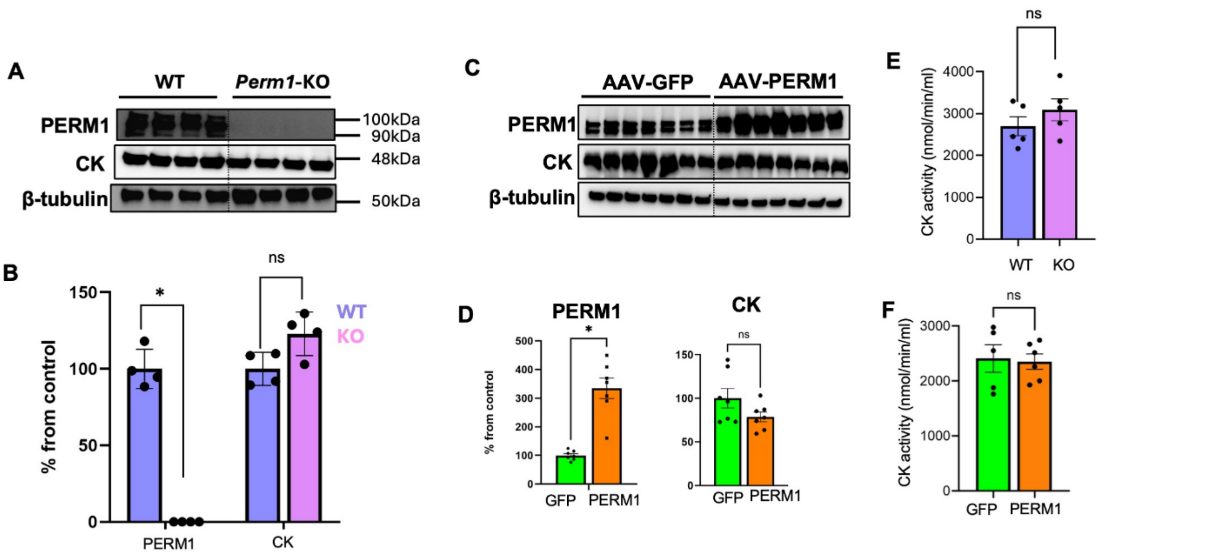


**Figure S3.** (**A-B**) Western blot analysis shows no change in creatine kinase expression in ***Perm1***-KO hearts. (**C-D**) AAV-mediated PERM1 overexpression does not alter creatine kinase expression levels. (**E-F**) Creatine kinase activity assays show no significant change in enzymatic activity in ***Perm1***-KO or AAV-PERM1 hearts. *Student's t*-test; $p < 0.05$; ns, not significant.

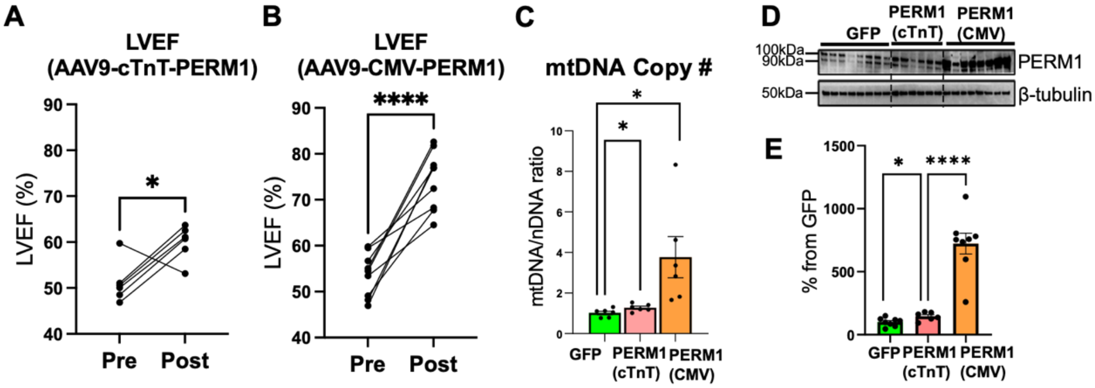


**Figure S4. Comparison of AAV-PERM1 vectors driven by a cardiac-specific promoter (cTnT) vs. a ubiquitous promoter (CMV) in C57BL/6N mice**. (**A-B**) Left ventricular ejection fraction (LVEF) shows a significant increase with both promoters, but a much stronger effect is observed with the CMV promoter. Similarly, the CMV promoter is superior to cTnT promoter in increasing mitochondrial DNA (mtDNA) copy number (**C**) and PERM1 expression (**D-E**). Paired t-test in **A** and **B**, one-way ANOVA in **C** and **E**. * $p < 0.05$; **** $p < 0.0001$.
